# Detecting copy number variations from single-cell chromatin sequencing data by AtaCNV

**DOI:** 10.1101/2023.10.15.562383

**Authors:** Xiaochen Wang, Zijie Jin, Yang Shi, Ruibin Xi

## Abstract

Singe-cell assay of transposase accessible chromatin sequencing (scATAC-seq) can unbiasedly profile genome-wide chromatin accessibility in single cells. In single-cell tumor studies, identification of normal cells or tumor clonal structures often rely on copy number variations (CNVs). However, CNV detection from scATAC-seq is difficult due to the high noise, sparsity, and confounding factors. Here, we describe AtaCNV, a computational algorithm that accurately detects high resolution CNVs from scATAC-seq data. We benchmark AtaCNV using simulation and real data and find AtaCNV’s superior performance. Analyses of 10 scATAC-seq datasets shows that AtaCNV could effectively distinguish malignant from non-malignant cells. In glioblastoma, endometrial and ovarian cancer samples, AtaCNV identifies subclones at distinct cellular states, suggesting important interplay between genetic and epigenetic plasticity. Some tumor subclones only differ in small-scale CNVs, demonstrating the importance of high-resolution CNV detection. These data show that AtaCNV can aid the integrative analysis for understanding the complex heterogeneity in cancer.

## Introduction

Assay of Transposase Accessible Chromatin sequencing (ATAC-seq)^1,2^ is widely used to measure the chromatin accessibility of whole genome. The development of single-cell ATAC-seq (scATAC-seq)^3^ further enables dissecting the epigenomic landscape and gene regulatory modules at single-cell resolution. scATAC-seq has been applied to a large number of tumor studies to discover epigenomic mechanisms for tumorigenesis and potential therapeutic approaches^4,5^. One important goal of single-cell tumor researches is to study the heterogeneity and evolution of tumor cells. Copy number variations (CNV), defined as gains or losses of DNA segments, are a major class of genomic variations in tumor cells. Different tumor subclones often have distinct CNV patterns, and CNVs may confer proliferation advantages and therapeutic resistance to cancer cells^6-8^. Accurate and comprehensive detection of CNVs is thus an essential step for single-cell tumor studies.

CNVs are usually detected by analyzing sequencing depths of genomic regions. Single-cell DNA sequencing (scDNA-seq) is probably the most suitable platform for single-cell CNV analyses, because its sequencing depths presumably are only affected by the copy numbers of genomic regions. However, CNV analyses are also often performed with other single-cell sequencing technologies^4,9^, such as single-cell RNA-sequencing (scRNA-seq) as well as scATAC-seq, to identify tumor subclones or to discern tumor and normal cells. Many CNV tools based on scDNA-seq and scRNA-seq data have been developed, such as SCOPE^10^, inferCNV^11^, and CopyKAT^12^. Recently, a few scATAC-seq CNV detection algorithms^13,14^ are also developed. However, these algorithms either suffer from limited detection resolution^13^ or require matched DNA sequencing data for breakpoint detection^14^.

Compared with scDNA-seq or scRNA-seq, CNV detection based on scATAC-seq is more challenging. (1) Unlike scDNA-seq, the sequencing depths of scATAC-seq are affected not only by CNVs but also by different chromatin accessibility across genomic regions. Accurate CNV detection based on scATAC-seq data requires proper normalization of scATAC-seq data to remove or reduce the influence of chromatin accessibility. (2) Compared with scRNA-seq data, scATAC-seq data are extremely sparse and noisy^15^. The numbers of CNVs and the positions of CNV breakpoints are usually unknown and must be estimated from the data. The high level of noise and sparsity can blur the difference between consecutive CNV regions and make detection and localization of CNV breakpoints very difficult.

To address these challenges, we develop a new CNV detection algorithm based on scATAC-seq data called AtaCNV. To account for the influence of chromatin accessibility on the sequencing depth, AtaCNV selects non-tumor cells and normalizes the sequencing depths against the non-tumor cells. The normalized sequencing depths of single cells are then jointly segmented^16^. The joint segmentation allows AtaCNV to pool information from neighboring genomic regions with the same CNV states and cells with similar CNV profiles, and thus could significantly denoise the data and make CNV signals more pronounced. Extensive simulation and real data benchmarking showed that AtaCNV can sensitively detect CNVs with high precisions. We further demonstrated the application of AtaCNV to scATAC-seq datasets from five different cancer types, including basal cell carcinoma (BCC), pancreatic ductal adenocarcinoma (PDAC), glioblastoma (GBM), ovarian cancer (OC), and endometrial cancer (EC), and revealed important subclones with distinct CNV profiles.

## Results

### Overview of the scATAC-seq CNV detection algorithm AtaCNV

AtaCNV mainly consists of a normalization step and a segmentation step (Figure 1A and **Methods**). After data pre-processing and quality control, AtaCNV generates a single-cell read count matrix over genomic bins of 1 million base pairs (1 Mbp). Firstly, cells and genomic bins are filtered according to bin mappability^17^ and number of zero entries. To reduce the extreme noisiness, AtaCNV then smooths the count matrix by fitting a one-order dynamic linear model for each cell. If normal cells are available, AtaCNV normalizes the smoothed count data against those of normal cells to deconvolute copy number signals from other confounding factors such as chromatin accessibility, mappability, and GC-content^17^. If normal cells are not available, observing that tumor single-cell data often contain many non-tumor cells, AtaCNV clusters the cells and identifies a group of high confidence normal cells and normalizes the data against their smoothed depth data. Finally, AtaCNV applies the multi-sample BIC-seq algorithm^16^ to jointly segment all single cells and estimates the copy number ratios for each cell in each segment. CNV burden scores are also derived and cells with high CNV scores are regarded as malignant cells (**Methods**).

**Figure 1.**
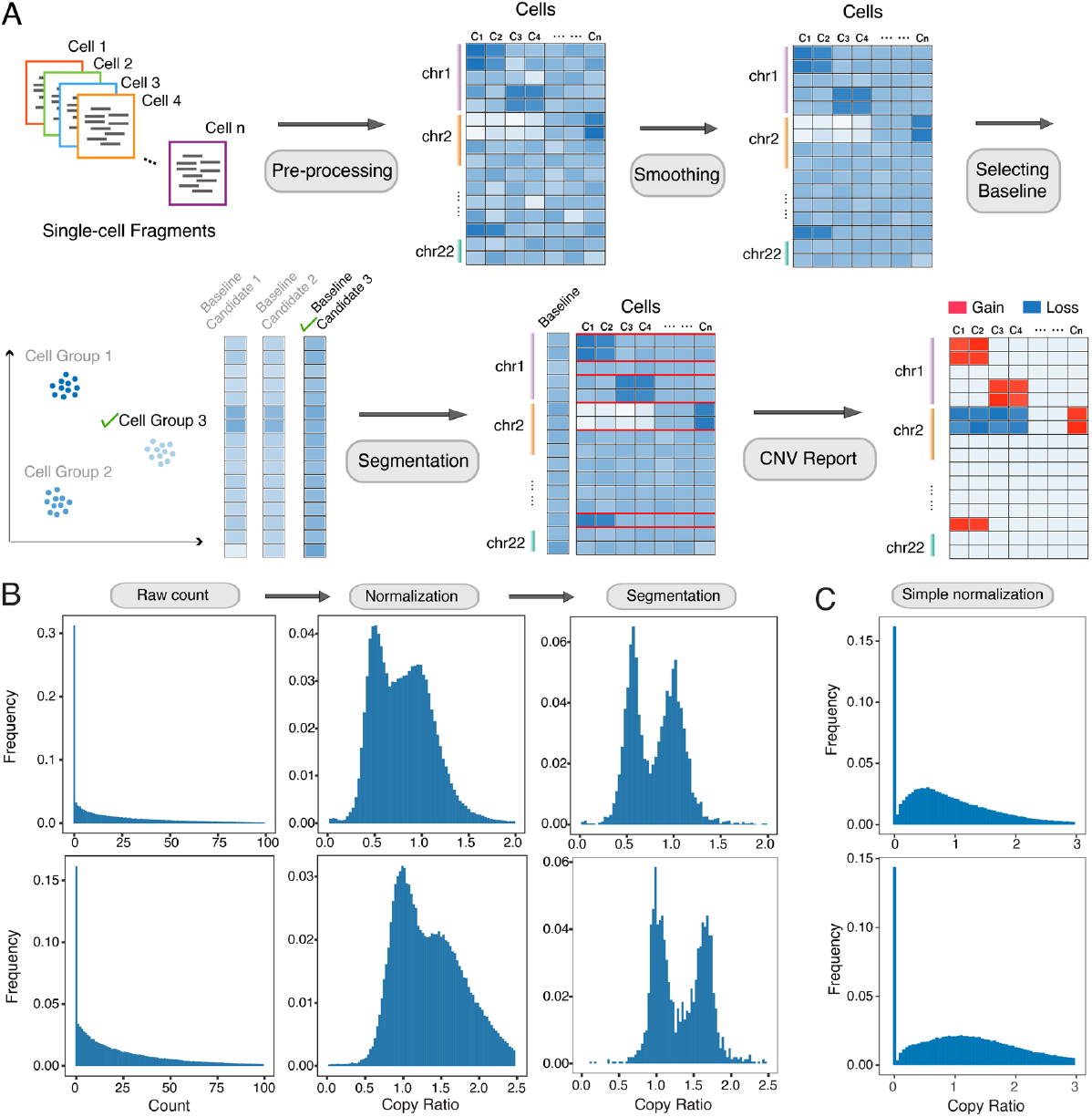
Overview of AtaCNV workflow. **(A)** AtaCNV takes read count matrix generated by scATAC-seq fragment file as input. In the normalization step, AtaCNV smooths the noisy read signals and defines a baseline from the preselected normal cells. Then AtaCNV performs segmentation and reports the final CNV profile. **(B)** The effect of the AtaCNV workflow on a GBM dataset. Distributions of the original read counts, copy ratios after normalization and copy ratios after segmentation in all cells are shown. The two rows in the plot correspond to the results in chromosome 10 and 7, respectively. **(C)** The distribution of copy ratios when applying a simple normalization method.

We applied AtaCNV to a GBM dataset^18^ and demonstrated the effect of the AtaCNV workflow (Figure 1B, C). It is well-known that GBM genomes often have CNVs at chromosome 7 and 10^19^. However, the raw counts of the bins at chromosome 7 and 10 do not show any peaks corresponding to different CNV states. After smoothing and normalization by the normal cells, CNV peaks start to occur, and after segmentation, CNV peaks become well separated (Figure 1B). In comparison, simple normalization by depths of normal cells cannot separate different copy number states (Figure 1C).

### AtaCNV detects CNVs with high sensitivity and precision in simulations

We compared the performance of AtaCNV with Copy-scAT and a simple baseline method using simulations (**Methods**). Briefly, simATAC^20^ was used to generate single-cell bin count matrices based on a user-specified reference scATAC-seq dataset. To simulate CNV events, we randomly chose genomic bins and altered their read depths by multiplying copy ratios to the underlying Poisson parameters. We considered two CNV-size setups, chromosome arm level CNVs and smaller scale CNVs of 10-20Mbp in length. The number of tumor subclones, noise levels, and overall coverage were also varied to evaluate the influences of these factors on the performances of these algorithms.

Overall, AtaCNV showed higher levels of precisions and recalls than the other two methods (Figure 2). In the arm-level simulations, the precisions and recalls of AtaCNV were over 0.9 and 0.75, respectively, and the overall F-scores were much higher (Figure 2A). In contrast, the recalls of the baseline method were extremely low, possibly because without proper normalization and segmentation, the CNV signals were buried in the biases and noises in the data. Copy-scAT had relatively high sensitivity but its precisions were much lower. This was possibly because in certain CNV regions, the Gaussian decomposition used by Copy-scAT misidentified which cells possessed the CNVs (Supplementary Figure 1). In the small scale CNV scenario, AtaCNV still showed a high level of recalls while keeping a similar level of precisions (Figure 2B). We also compared these methods in terms of their accuracies in estimating the copy ratios and found that AtaCNV outperformed others (Supplementary Figure 2). In addition, we found that the performance of AtaCNV remained largely the same with different sequencing coverages, noise levels and numbers of tumor subclones.

**Figure 2.**
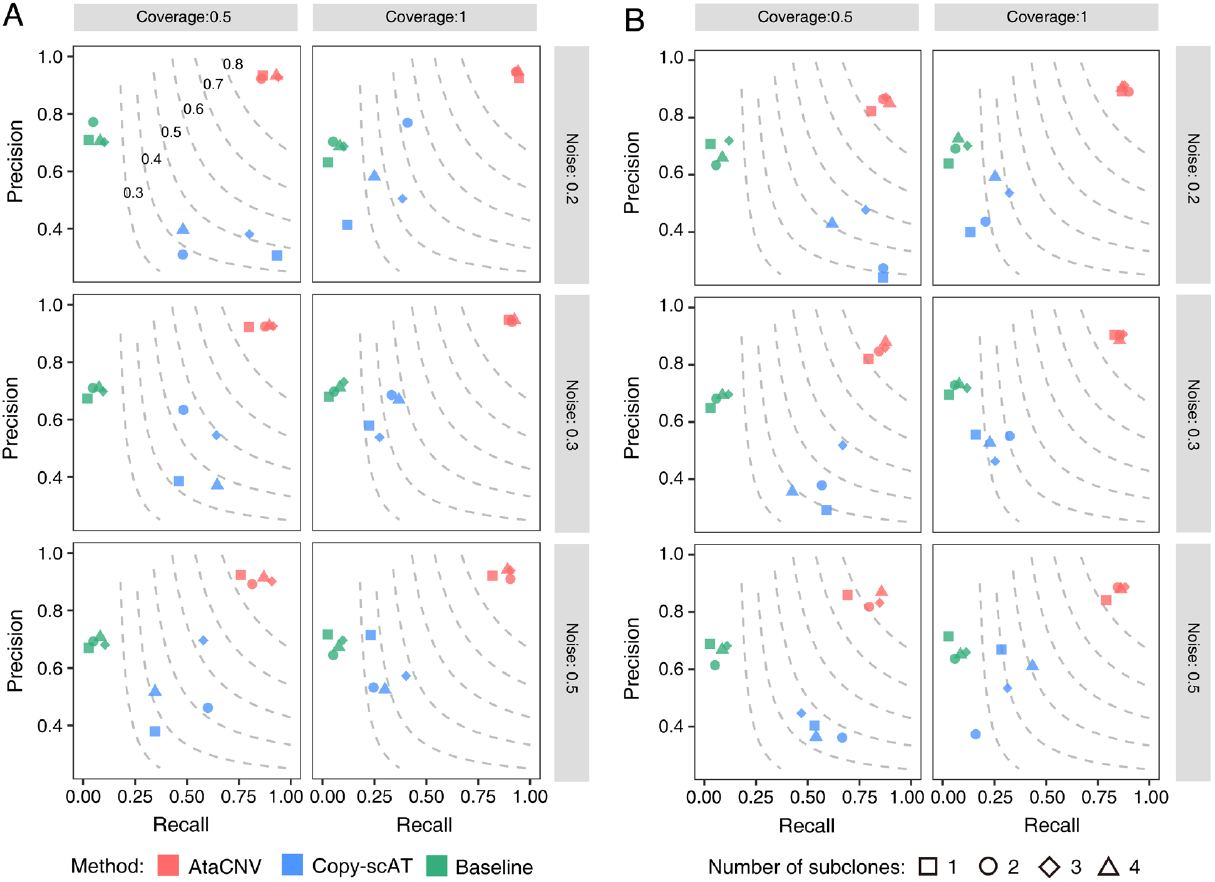
Performance comparison using simulation. **(A, B)** The precisions and recalls of AtaCNV, Copy-scAT, and the baseline method in arm-level CNV simulations (A) and small-scale CNV simulations (B). The dashed lines are contour lines with constant F-scores (F-scores are marked in the top-left figure).

### Benchmarking with a BCC scATAC-seq dataset

We further tested the ability of AtaCNV using real scATAC-Seq datasets profiled from a patient with BCC^4^. The datasets contain BCC tumor samples and paired normal sample sequenced before and after immunothserapy. CNV results of the post-treatment sample are displayed in Figure 3. Among all 309 cells from the tumor sample, 115 were annotated as tumor cells, while the remaining were annotated as non-tumor cell types such as myeloid and fibroblasts^4^. Matched whole-exome-sequencing (WES) data was also available^21^ and we compared CNVs inferred from scATAC-seq with those from the WES data (Figure 3A-E). AtaCNV was able to detect most CNVs called from the WES data, such as amplifications of chromosome 3q, 6p, 8, 20, and 21, and deletions of chromosome 17 and 22. In comparison, Copy-scAT only reported amplifications of chromosome 3q and 8q. In addition, Copy-scAT falsely reported amplifications of chromosome 8q in 39 normal cells (Figure 3B). We further evaluated the performances of two methods with or without reference data of known normal cells. The reference data was chosen as either the annotated normal cells or the paired normal cells. With different choices of the reference data, CNVs detected by AtaCNV were more similar to CNVs from the WES data than by Copy-scAT (Figure 3F, G), as measured by correlation or absolute difference with the WES log2 copy ratios (**Methods**).

**Figure 3.**
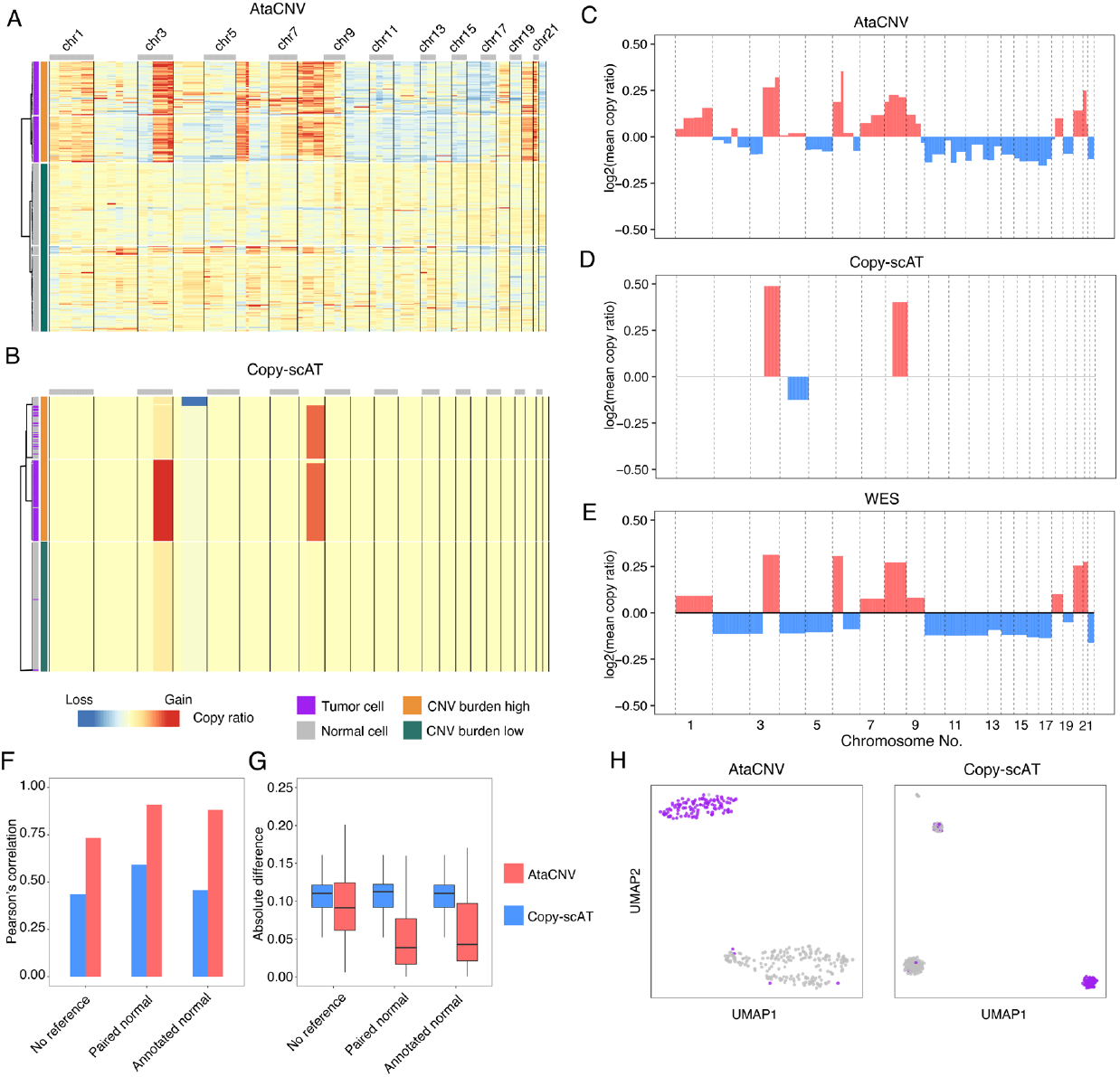
Performance evaluation using a BCC sample. **(A, B)** The CNV profiles of the SU008 BCC sample reported by AtaCNV (A) and Copy-scAT (B) using the paired normal samples as reference. Tumor cells indicated by purple bar and normal cells indicated by grey bar were all from the BCC tumor sample and annotated in the original research. **(C)** Mean copy ratios over all cells of genomic segments given by AtaCNV. **(D)** Mean copy ratios over all cells of genomic segments given by Copy-scAT. **(E)** The copy ratio profile inferred by matched WES dataset. **(F)** The Pearson’s correlations between WES copy ratios and mean copy ratios given by AtaCNV and Copy-scAT using different reference data. **(G)** The absolute differences between WES copy ratios and mean copy ratios given by AtaCNV and Copy-scAT. **(H)** The UMAP plots of CNV profiles from AtaCNV and Copy-scAT. Points are colored by their tumor/normal labels.

We next evaluated the ability of AtaCNV in distinguishing tumor cells from non-tumor cells. Hierarchical clustering (Figure 3A) and UMAP projection (Figure 3H) of the CNV profiles given by AtaCNV showed two distinct cell groups. As expected, the cell group with a high CNV burden corresponded to the annotated tumor cells, and the group without much CNVs corresponded to the annotated non-tumor cells. In contrast, Copy-scAT was less accurate in separating the two cell types (Figure 3B, H). In the pre-treatment sample from the same patient, AtaCNV also performed similarly (Supplementary Figure 3A-E).

### AtaCNV accurately distinguished tumor and normal cells in various cancer types

In the analysis of single-cell sequencing data from tumor samples, it is common to use CNV to distinguish tumor cells. Therefore, we further validated the effectiveness of AtaCNV in this regard. We applied AtaCNV to multiple types of tumor scATAC-seq data, including two PDAC datasets^22^, four adult GBM (aGBM) datasets^18^, two pediatric GBM (pGBM) datasets^13^, one OC dataset and one EC dataset^23^. In all datasets, we performed cell clustering using ArchR^24^ and manually annotated tumor cell clusters based on the predicted expression scores of tumor marker genes (**Methods**). The total numbers of cells and tumor cells in each dataset is shown in Figure 4A. Other summary statistics of the datasets are shown in Supplementary Table 1. We obtained single-cell level CNV results using AtaCNV and Copy-scAT, respectively, and subsequently identified tumor cell clusters enriched with CNVs (**Methods**). AtaCNV achieved higher accuracies in distinguishing tumor cells than Copy-scAT (Figure 4B), where accuracy is defined as the number of correctly classified cells divided by the total number of cells. We also calculated the CNV burden at single-cell level and examined the distribution of CNV burden in normal cells versus tumor cells. The median CNV burdens of tumor cells given by AtaCNV were consistently, often significantly, larger than those of normal cells for all the datasets (Figure 4C). In contrast, CNV burdens of tumor cells given by Copy-scAT were smaller than those of normal cells in some datasets such as aGBM 4218 and OC 3CCF1L (Figure 4D), explaining Copy-scAT’s inferior performance in distinguishing tumor and normal cells in these datasets (Figure 4B).

**Figure 4.**
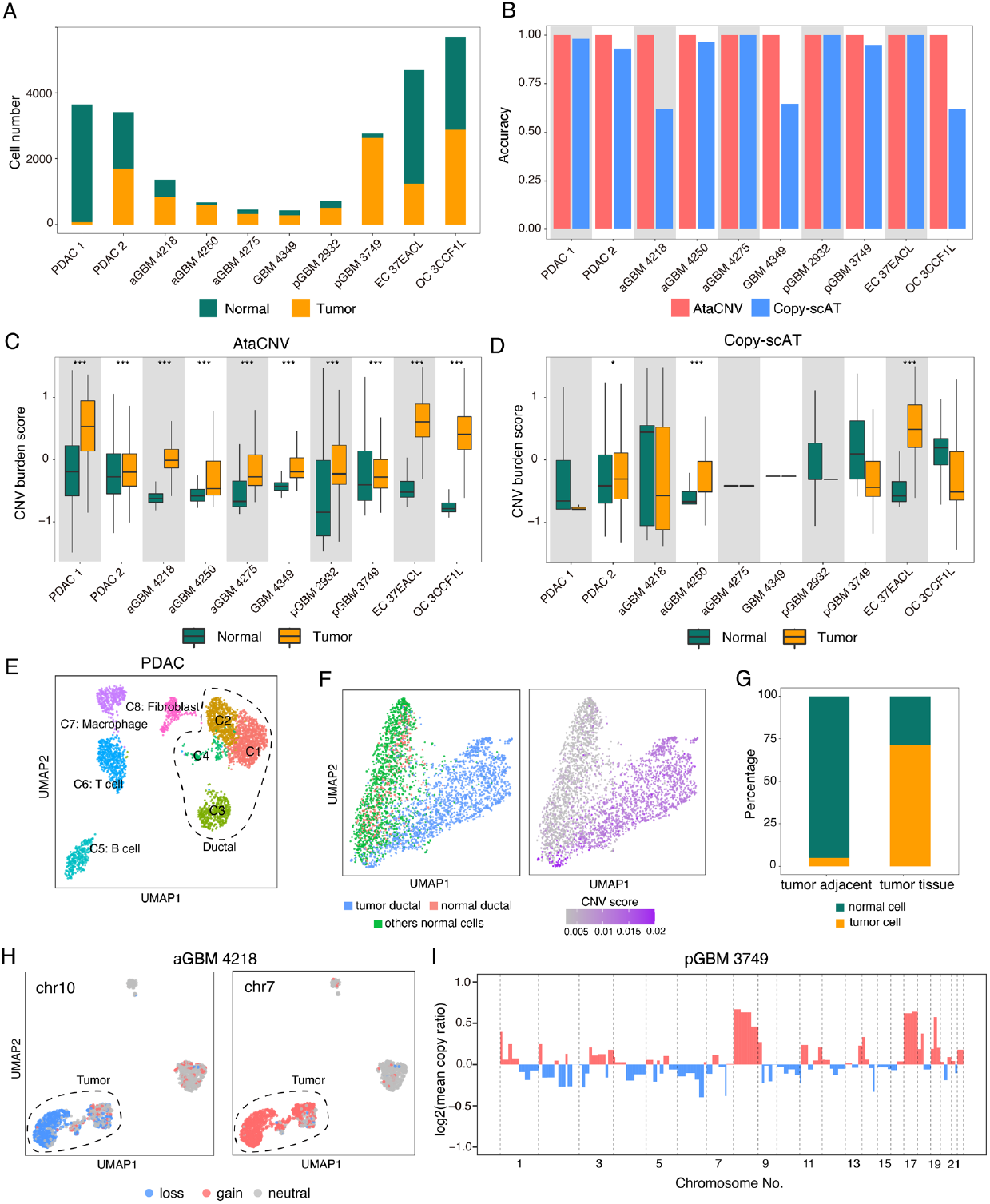
Tumor cell classification in various cancer types. **(A)** Barplots of tumor and normal cell numbers. Tumor cells are manually annotated using tumor marker genes **(B)** Barplots of classification accuracies of AtaCNV and Copy-scAT. **(C)** Boxplots of CNV scores of annotated tumor and normal cells given by AtaCNV. The asterisks indicate that the scores of the tumor cells are significantly higher than those of the normal cells (one-sided Wilcoxon’s rank sum test^40^. * *P* < 0.05; *** *P* < 0.001). **(D)** Boxplots of CNV scores of annotated tumor and normal cells given by Copy-scAT. **(E)** The UMAP plot of PDAC cells based on peak signal data. **(F)** The UMAP plots of PDAC cells using the CNV profile predicted by AtaCNV. The left plot shows the classification of tumor/normal ductal cells and other normal cells. The right plot shows the CNV scores of each cell. **(G)** The barplots of proportions of malignant cells in PDAC tumor and tumor adjacent tissues. The proportions are estimated based on AtaCNV’s classification. **(H)** The UMAP plots of aGBM 4218 cells colored by copy number states of chromosome 7 and 10. **(I)** The inferred mean copy ratio profile of pGBM 3749 by AtaCNV.

We further demonstrated the effectiveness of AtaCNV in a PDAC dataset. This dataset contains a PDAC tumor sample (n=1149 cells) and its adjacent pancreatic para-tumor tissue (n=2313 cells). The clustering results yielded 8 cell clusters, which were annotated as T cells, B cells, macrophages, fibroblasts and ductal cells using marker genes (Figure 4E and Supplementary Figure 4A-C). We employed transcription factor enrichment analysis (**Methods**) within ductal cells to identify tumor cells and observed that C1, C2 and C4 were more enriched with many proto-oncogenes (such as *JUN, JUNB, JUND* from the JUN family and *FOSL1, FOSL2, FOS, FOSB* from the FOS family, Supplementary Figure 4D). These tumor cells were also identified by AtaCNV. Tumor ductal cells were readily separated from normal ductal cells in the UMAP space of the inferred copy ratios and evidently had higher CNV burden (Figure 4F). The coexistence of the two types of ductal cells indicated that tumor cells may evolve from normal ductal cells^25^. Furthermore, as expected, tumor tissue contained much more tumor cells than the pancreatic para-tumor tissue (Figure 4G and Supplementary Figure 4E).

Using scATAC-seq data from adult and pediatric GBM patients, AtaCNV was also validated by copy numbers in specific genomic regions. AtaCNV classified 839, 577, 314, and 279 cells in four aGBM samples as tumor cells, respectively, and classified 502 and 2,628 tumor cells in two pGBM samples. The classification results were validated by the expression of the marker gene *EGFR* (Supplementary Figure 5). It is well-known that amplifications of chromosome 7 and deletions of chromosome 10 occur highly frequently in aGBMs but not so much in pGBMs^26^. Among four aGBM samples, most of the aGBM tumor cells harbored both gains of chromosome 7 and losses of chromosome 10 (Figure 4H and Supplementary Figure 6A-C). Meanwhile, these gains or losses occurred much less frequently in tumor cells of pGBM 3749 and 2932 (Figure 4I and Supplementary Figure 6D). We also applied Copy-scAT and found that it could not accurately capture these CNVs in aGBMs (Supplementary Figure 6E). These results demonstrated that AtaCNV was capable of distinguishing tumor cells in various cancer types and accurately detecting CNVs in tumor cells.

### AtaCNV provides mechanistic insights of intra-tumor heterogeneity in adult GBM

We further analyzed the sub-clonal structure of aGBM 4218. The CNV profile revealed three distinct tumor clusters, S1, S2, and S3 (Figure 5A, B). Subclone S1 had a unique strong amplification of chromosome 8, but did not have the strong amplifications of chromosome 7 that were detected in subclone S2 and S3. Subclone S1 and S2 had a shared amplification of chromosome 18q that S3 did not have (Figure 5C). Overall, similarities between the copy number profiles of S2 and S1 (Pearson’s correlation=0.74) or S2 and S3 (Pearson’s correlation=0.76) are much higher than between S1 and S3 (Pearson’s correlation=0.54). Occurrence of CNVs on genomic segments suggested that S2 harbored very few unique CNVs that S1 and S3 did not have (Figure 5D). These CNV features indicated that S2 might represent an intermediate state between S1 and S3.

**Figure 5.**
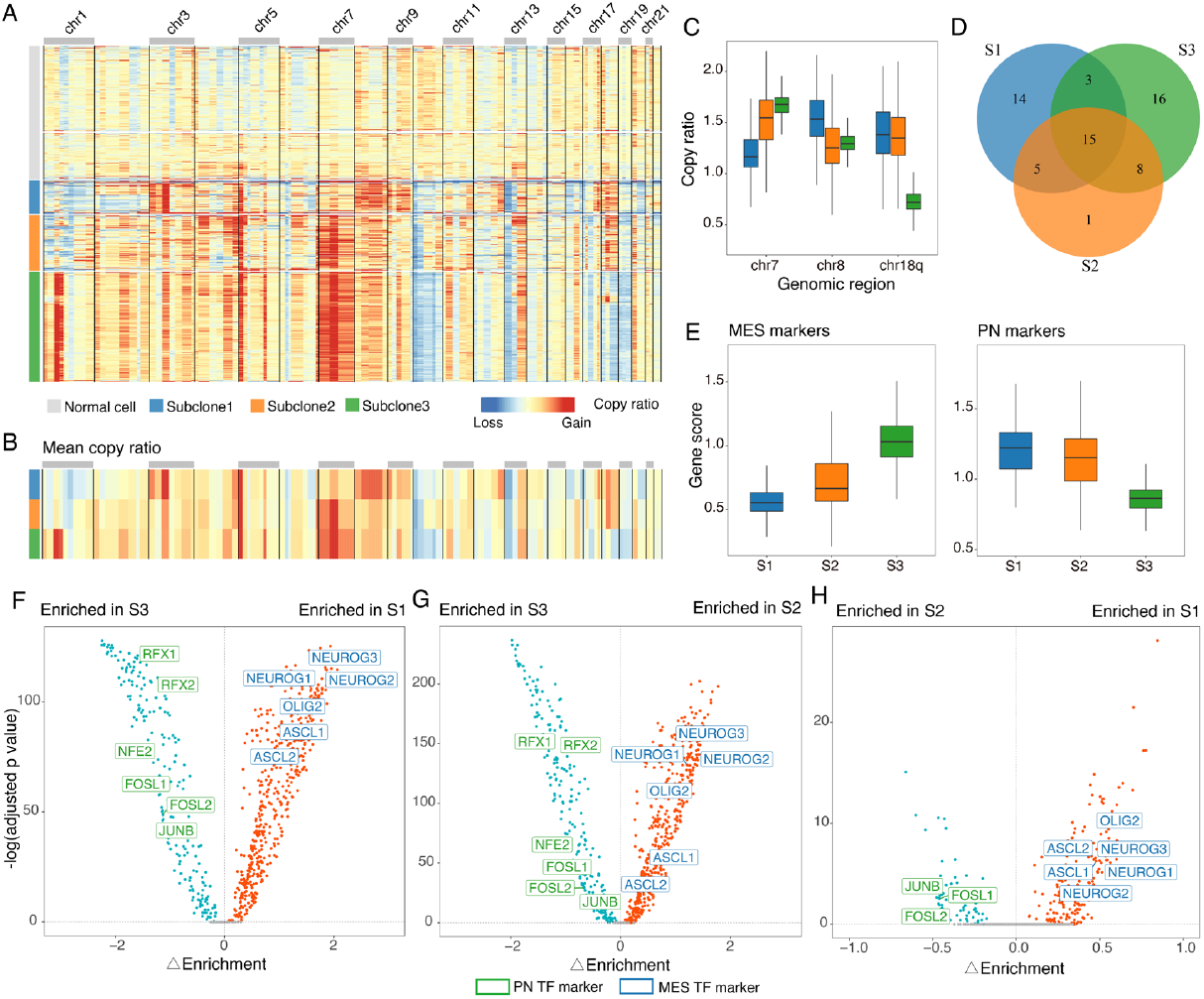
Subclone analysis of aGBM 4218. **(A)** The CNV profile reported by AtaCNV. The left bars indicate the normal cells and three tumor subclones estimated based on the CNV profile. **(B)** Mean copy ratios of cells in the three tumor subclones. **(C)** Boxplots showing copy ratios of several genomic regions in the three subclones. **(D)** Venn plot showing the numbers of genomic segments with CNVs in the three subclones. **(E)** Boxplots of MES and PN marker gene scores in the three subclones. **(F-H)** The differentially enriched TF motif tests between S3 and S1 (F), between S3 and S2 (G), and between S2 and S1 (H). The x-axis is the enrichment difference and the y-axis is the negative log adjusted p-values.

Previous single-cell transcriptomic analysis showed that aGBMs might contain cells of different subtypes including proneural (PN), neural (NL), mesenchymal (MES), and classic (CL) subtypes^9^, and that aGBM cells could be described in a single axis ranging from PN to CL and then to MES^27^. We hypothesized that S1, S2 and S3 might correspond to these known aGBM cell subtypes. In fact, using previously obtained marker genes of PN and MES from scRNA-seq^27^, we found that cells in S1 and S3 had higher mean expression scores of PN and MES markers, respectively, while S2 always lied in the middle level (Figure 5E). From marker genes of S1-S3 identified solely using scATAC-seq (**Methods**), we also found that S3 was enriched with MES markers (*CD44, EGFR, CHI3L1*), S1 was enriched with PN markers (*ASCL1, DLL3* and *OLIG2*) and S2 was in an intermediate state (Supplementary Figure 7). We then performed pairwise differential motif enrichment tests to identify related transcription factors (**Methods**). From the differentially enriched TFs between S1 and S3, we could see that MES TF markers and PN markers were enriched in S1 and S3, respectively (Figure 5F), which again validated the identities of these two populations. However, when comparing S2 to MES subclone, PN markers were significantly enriched (Figure 5G). Similarly, when comparing S2 to PN subclone, MES markers were significantly enriched (Figure 5H). Collectively, consistent with the CNV profiles, these results all supported that S2 might be an intermediate population between S1 and S3.

### AtaCNV identifies subclones with distinct transcriptomic features in human gynecologic cancers

We applied AtaCNV to study subclones of two human gynecologic cancers, including one OC sample and one EC sample^23^. Previous study^23^ assigned histological categories for the samples, where the former was classified as ovarian carcinosarcoma and the latter as serous EC. In both samples, we identified tumor cells based on the CNV profile, which could be validated by cancer marker gene *MUC16* and *WFDC2* (Supplementary Figure 8). In the OC sample, AtaCNV detected recurrent broad CNVs of ovarian tumor^28^ including copy number gain of chromosome 3q, 8q, 20 and copy number loss of chromosome 17p, 22 (Figure 6A). The CNV profile suggested that there existed two subclones within tumor cells (Figure 6A and 6B). We used Seurat^29^ to integrate the scATAC-seq dataset with the simultaneously sequenced scRNA-seq data^23^ (**Methods**) and investigated the transcriptomic feature of these two subclones. We applied gene set variation analysis (GSVA)^30^ at the single-cell level to identify differentially enriched cancer hallmark pathways (Figure 6C and **Methods**). Subclone S1 showed significantly higher epithelial-to-mesenchymal transition (EMT) score than subclone S2, while subclone S2 showed higher score of apical junctions, a hallmark of polarized epithelial cells. EMT is a key process in which epithelial cells lose polarity and develop mesenchymal features. Its role in ovarian cancer has been extensively studied^31,32^. In carcinosarcoma, researches suggested that a higher EMT score corresponded to the sarcoma component while a lower EMT score corresponded to the carcinoma component^31,33^. Therefore, S1 and S2 were likely to be comprised of sarcoma and carcinoma component, respectively. This assumption was also supported by the expression of epithelial marker genes and mesenchymal marker genes in two subclones (Figure 6D, E).

**Figure 6.**
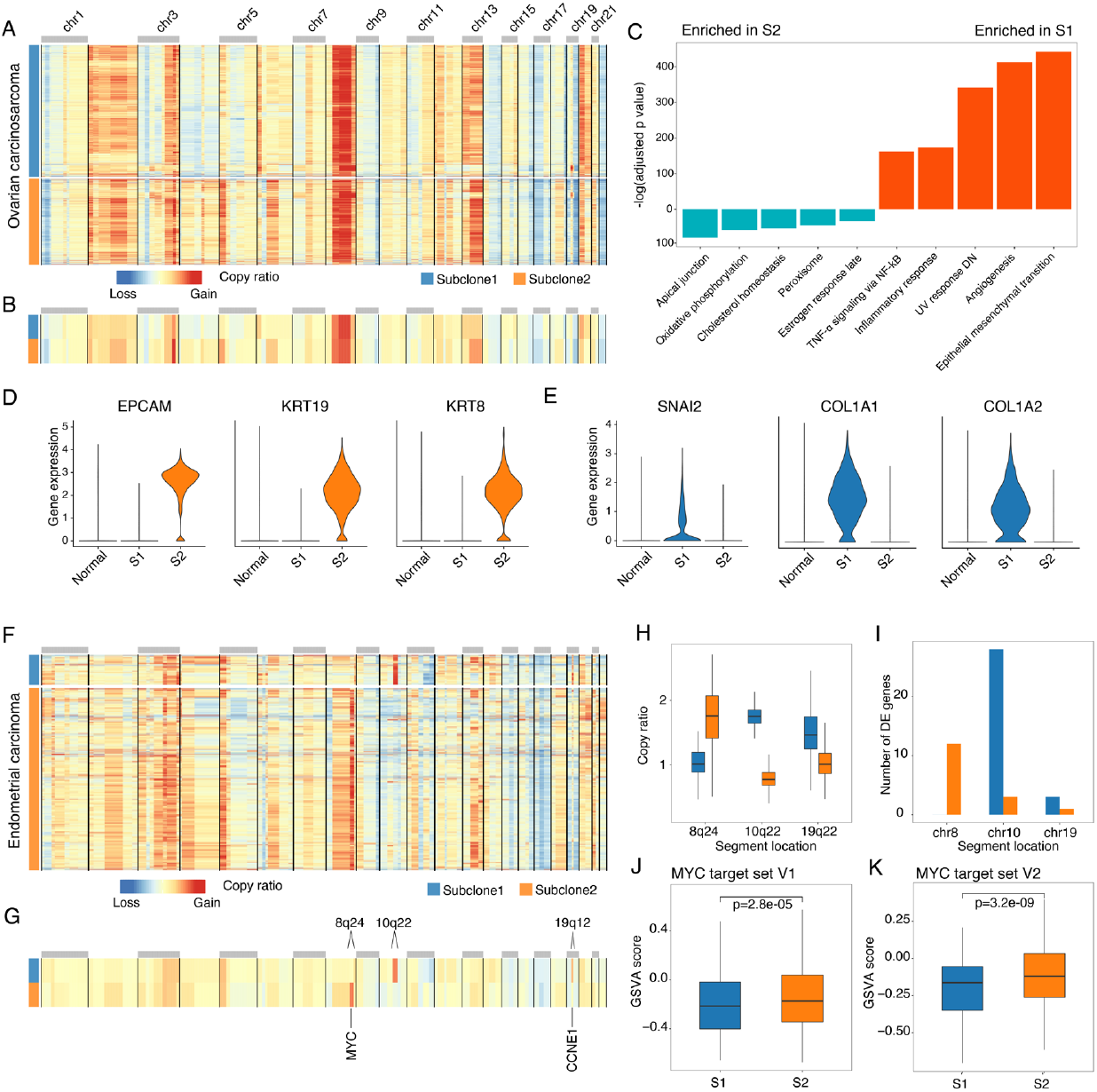
Subclone analyses of human gynecologic cancers. **(A)** The CNV profile of tumor cells in OC 3CCF1L reported by AtaCNV and the two inferred tumor subclones. **(B)** Mean copy ratios in the two subclones of OC 3CCF1L. **(C)** Differential enriched cancer hallmark pathways between the two subclones of OC 3CCF1L. **(D, E)** Violin plots of epithelial (D) and mesenchymal (E) marker gene expressions in the normal and two tumor subclones of OC 3CCF1L. **(F)** The CNV profile of tumor cells in EC 37EACL reported by AtaCNV and the two inferred subclones. **(G)** Mean copy ratios in the two subclones of EC 37EACL. **(H)** Boxplots of copy ratios of three genomic segments in the two subclones. **(I)** Numbers of differentially expressed genes in these segments. Bars of different colors show the numbers of up-regulated genes in the corresponding subclone. **(J, K)** Boxplots of GSVA scores of MYC target set V1 (J) and V2 (K) in the two subclone cells. The p-values marked in the plots are obtained by the Wilcoxon’s rank sum test.

In the EC sample, AtaCNV also detected two subclones with distinct CNV profiles (Figure 6F, G). Previous study classified ECs into four categories: POLE ultra-mutated, MSI hyper-mutated, copy number low, and copy number high^34^. The frequent CNV events indicated that this sample belonged to the copy-number high subtype, which was a major type of serous carcinoma. The two subclones differed in copy numbers of several relatively small genomic segments (Figure 6F and 6G), where the three most distinct CNVs were clone-specific (Figure 6H). This was supported by the paired scRNA-seq data: genes in these regions tended to have higher expression in the corresponding subclone (Figure 6I). Subclone 1 was characterized by focal amplification of two segments that overlapped with chromosome 10q22 and 19q12, respectively. Both amplifications were reported to be frequently occurred in EC^35^ and the latter harbored the oncogene *CCNE1*. Subclone 2 had a focal amplification of a segment overlapping with 8q24, which is recurrently amplified in EC^35,36^. The amplification contains the gene *MYC*, whose oncogenic role in ECs had been discussed^34^. GSVA analysis of the paired scRNA-seq data showed that the scores of *MYC* target gene sets were significantly higher in S2 than in S1 (Figure 6J, K). These results suggested that subclones identified by AtaCNV were bona fide subclones and might have distinct phenotypic features. Meanwhile, Copy-scAT could only detect the focal amplification at 10q22 (Supplementary Figure 9). This application revealed the advantage of AtaCNV in detecting focal CNVs, which is necessary for accurate subclone analyses.

## Discussion

In this paper, we present AtaCNV, a CNV detection algorithm based on scATAC-seq data. CNV detection using scATAC-seq data is challenging because of the high noise, high sparsity, and the existence of confounding factors in the data. By local smoothing, normalization against known or proper selected normal cells and joint segmentation, AtaCNV achieves precise arm-level and small-scale CNV detection without requiring a matched DNA sequencing data, as demonstrated by our extensive simulation and real data analyses. The CNV profile given by AtaCNV can be used to distinguish tumor and normal cells and to analyze tumor heterogeneity.

We demonstrated applications of AtaCNV for tumor heterogeneity analyses. In the GBM dataset, we identified three subclones based single-cell CNV profiles. The three subclones had distinct expression patterns. S1 highly expressed the PN-subtype genes, S3 expressed MES-subtype genes, and S2 was in an intermediate state between S1 and S3. This corresponds with the single-axis explanation of GBM subtypes. Recent findings also classified GBM cells into AC-like, MES-like, OPC-like and NPC-like cells based on gene expression profiles^37^. *EGFR* is a key oncogene on chromosome 7, with the highest expression in AC-like cells and the lowest in OPC-like cells. In this sample, we can clearly observe that chromosome 7 is not amplified in S1, which is consistent with the fact that OPC-like cells mainly occur in the PN subtype. This further demonstrates the impact of copy number variations on transcriptomic events. In the EC dataset, AtaCNV identified two subclones that mainly differed in three focal CNVs (8q24, 10q22 and 19q12). In the GBM and OC datasets, other than large arm-level CNVs, different subclones also differed in focal CNVs (e.g. 1q32 in GBM, 3q26 in OC). These focal CNVs often contained important cancer genes (e.g. *MYC* at 8q24, *SOX2* at 3q26) that might play important roles in the survival and proliferation of the tumor subclones. Without accurate high-resolution CNV detection as given by AtaCNV, identification of these subclones would be difficult or even impossible.

Recently, improvements in single-cell sequencing protocols have enabled simultaneous sequencing of multiple omics. Chromatin accessibility and gene expression can be simultaneously profiled ^38,39^. scRNA-seq data have higher read depth but the confounding problem (e.g. by gene expression) might be more prominent. scATAC-seq data directly profile DNA sequences, but the signals are weaker and sparser than scRNA-seq. CNV detection could be further improved by integrating both types of data. In fact, we analyzed paired scATAC-seq and scRNA-seq data from gynecologic cancers, and found that CNV profiles inferred by two types of data were largely consistent (Supplementary Figure 10), suggesting that multi-omics CNV analysis is feasible and might render more robust results. Overall, our results strongly validate the performance of AtaCNV on scATAC-seq data. We expect that AtaCNV will have wide applications in single-cell tumor studies involving scATAC-seq data.

## Methods

### The AtaCNV method

#### Preprocessing

For each tumor sample, we first generate fragment files using Cell Ranger ATAC and perform quality control by the R package ArchR^24^. By default, only barcodes with *unique_fragments* > 3000 and *TSS_enrichment_score* > 4 are kept as cellular barcodes. To remove potential doublets, we also calculate the doublet score using ArchR’s function and filter out barcodes with doublet enrichment scores > 1.

We separate each chromosome into non-overlap bins of 1 million bp (Mbp). We generate a bin-by-cell read count matrix from the fragment file. To remove regions with low mappability, bins that contain few uniquely mappable positions (< 50%) are removed. All the columns and rows with over 80% zeroes will also be removed. We get a *m* × *n* count matrix ***X*** (*m* bins and *n* cells) in this step.

#### Normalization

The count matrix ***X*** is extremely noisy. To smooth the noisy count signals, we transform ***X*** using log(***X*** + 1) and fit a simple one-order dynamic linear model to each column of ***X***:

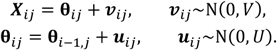

The model estimation is realized by R package dlm^41^. We empirically set *V* = 0.3 and *U* = 0.05 by default. Then, we replace ***X***_*ij*_ by the estimated **θ**_*ij*_ and still denote this matrix by ***X***. The smoothed matrix represents the denoised read depth of each cell in each bin.

Besides the copy number, read depth might be affected by various factors. We first convert ***X*** to count per million to remove the effect of library size. We assume that

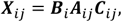

where ***B***_*i*_ represents the bin specific factors like GC content and mappability, ***A***_*ij*_ represents the different chromatin accessibility of each cell in each bin and ***C***_*ij*_ represents true copy number.

The factor ***A***_*ij*_ is mainly driven by different cell types that might carry different chromatin accessibility in the same region. However, such difference depends on the bin length. In non-tumor samples, the average read depths in 1 million bp length bins shows little difference between cell types (Supplementary Figure 11). Therefore, we may assume that ***A***_*ij*_ is also a bin specific factor.

To remove bin specific bias and deconvolute the true copy ratio, we need to estimate a read depth baseline of non-tumor cells. Since most tumor samples contain a certain percent of normal cells, we aim to divide cells into several clusters and preselect clusters of normal cells. Here, we apply ArchR’s clustering method. The method uses count matrix of 500 bp bins that represents peak level signals and applies an iterative Latent Semantic Indexing (LSI) algorithm to do single-cell clustering.

If we already know which cluster consists of non-tumor cells from additional information, we will subset these cells from ***X*** and use median value of each row as baseline. We denote this m-length vector as ***x***^*ref*^. However, such information is not directly available in most cases. In AtaCNV’s pipeline, we provide two strategies for normal cluster selection and their effects are shown in Supplementary Figure 12.

When scATAC-seq data of paired non-tumor sample is available, we preprocess it in the same way and get a smoothed read depth matrix ***X***^*paired*^. Then, the row’s medians of ***X***^*paired*^ could be used as a baseline. Denote this *m*-length vector as ***x***^*paired*^. To further rectify the difference between samples, we attempt to find a subset of normal cells from the original tumor sample instead of using ***x***^*paired*^ directly. For each cluster of the tumor data, the row’s median of the corresponding submatrix of ***X*** is compared with ***x***^*paired*^ using Pearson’s correlation. Clusters with correlations larger than 0.95 and having more than 20 cells were viewed as normal cells in the tumor sample. Then median read depths of these normal cells are used as the baseline (denote as ***x***^*ref*^). If there is no such group, we directly use ***x***^*paired*^ as the baseline.

When non-tumor data is unavailable, we propose to use the following method to select the normal clusters. Though the chromatin accessibility is not completely decided by gene transcription events, we observe that the read depth is correlated with the number of genes in the bin (Supplementary Figure 12A). In tumor clusters, such correlation is weakened by the influence of CNVs. Denote the numbers of genes in *m* bins by a *m*-length vector ***g***, we use ***g*** as a pseudo baseline and choose the cluster with the highest Pearson correlation with ***g*** as the normal cells. The baseline ***x***^*ref*^ is then calculated from these inferred normal cells.

After obtaining the baseline vector ***x***^*ref*^, we get the copy ratios for the *j*-th cell in the *i*-th bin as:

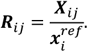

### Segmentation

To further denoise the data and detect the potential CNV boundaries, we use the multi-sample BIC-seq algorithm for segmentation. Based on the bin read counts of tumor and normal genomes, the algorithm uses the BIC criterion to merge bins iteratively and report the final segmentation result. Here, we use BIC-seq to determine common break points of all cells from the read depth matrix ***X*** and baseline ***x***^*ref*^.

Specifically, we denote the segmentation of *m* bins as a vector *S* = (*s*_0_, *s*_1_, …, *s*_*L*−1_, *s*_*L*_), where *s*_0_ = 0 < *s*_1_ < ⋯ < *s*_*L*−1_ < *s*_*L*_ = *m* represent the positions of breakpoints and the *l*-th segment consists of consecutive bins from *s*_*l*−1_ + 1 to *s*_*l*_. We minimize the following objective function:

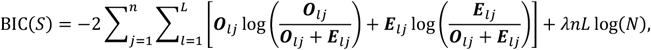

where 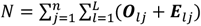 represents the total observed read depths of *j*-th cell in *l* -th segment and ***E***_*lj*_ represents the total expected read depths. They are calculated from ***X*** and ***x***^*ref*^, respectively:

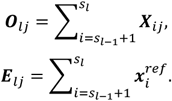

Tuning parameter *λ* is set to 5 by default. After obtaining the optimal segmentation 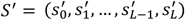, copy ratio of bins in the *l*-th segment is recalculated as:

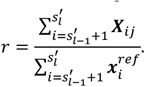

We denote the final result as a *m* × *n* copy ratio matrix ***R***.

### Simulation study

We utilized the R package simATAC^20^ to generate single-cell bin count matrix with CNV events. simATAC learns features from a given bin-by-cell count matrix and simulates similar read count matrices. We chose the bin length of the reference read count matrix as 1 Mbp, since both AtaCNV and Copy-scAT uses 1 Mbp non-overlapped bins as inputs. For each entry, simATAC generates read counts from a Poisson distribution, where the parameter of the Poisson distribution is learned from the reference dataset. Then, we multiply the copy ratio to the Poisson mean to generate bin counts in specific CNV regions. We further multiply a common coefficient to simulate counts with various overall coverage. Besides, simATAC can add Gaussian noises to the final count, which could be used to simulate different noise level. Details of the simulation are given below.

We first generated a bin-by-cell copy number matrix ***C***. We divided 1000 cells into five equal-sized groups and chose *m* groups to be tumor cell groups. For cells in a given tumor group, in chromosome arm-level CNV simulations, we randomly chose 5 chromosome arms for each tumor group. For each chosen arm, copy numbers of cells in the group were randomly sampled from {0,1,3,4} to simulate different aneuploid states. In small-scale CNV simulations, in addition to randomly selecting 4 chromosome arms, we also randomly chose 10∼20 consecutive bins on 4 different chromosomes to represent CNV segments. For the selected 4 arms and 4 segments, copy numbers of cells in the group were also randomly sampled from {0,1,3,4}. All other elements of ***C*** were set to be 2, which represent diploid state.

We then generated a matrix ***X*** of the same size of ***C*** that represented scATAC-seq read counts corresponding to the given copy number states ***C***. We used a 10X PBMC dataset as the reference data for simulation. From the reference, simATAC estimated a Poisson distribution Poisson(***λ***_*ij*_) for each entry *X*_*ij*_. To simulate copy number states, we varied parameter *sparse*.*fac* in simATAC, which was a factor multiplied to the parameter of the

Poisson distribution before simulating the final counts. We set this parameter to 0.5α to simulate counts with copy number *C*. Here, *α* controlled the overall coverage. Finally, we set mean and standard deviation of Gaussian noise to be -0.3 (the recommended value by simATAC) and *σ*, respectively. The whole process could be summarized as:

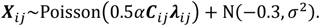

In general, *m* controls malignancy, *α* controls the overall sparsity and σ controls noise level. We varied *m* from {1, 2, 3, 4}, α from {0.5, 0.75, 1} and σ from {0.2, 0.3, 0.5}. For each setting, we simulated arm-level CNV and small-scale CNV for five repetitions.

The simulated bin count matrices were directly used as inputs of AtaCNV and Copy-scAT. We ran Copy-scAT using its default parameters. For both methods, we provided 200 non-tumor cells as reference data. We also added a baseline method for comparison, in which we divided read counts by the median of normal cells to calculate copy ratios directly.

We evaluated the performance of these methods by comparing the output copy ratio matrix ***R*** (Copy-scAT reported the copy numbers of chromosome arms and focal amplifications in specific regions. We also turned it into a copy ratio matrix.) to the ground truth copy number ***C***. Elements of ***R*** larger than 1.25 were defined as gains and those less than 0.75 were defined as losses. The others were defined as neural. The ground truth copy states of gain, loss, or neural were obtained from ***C***. Then we could calculate accuracy as well as precision and recall. We also calculated the mean square error (MSE) by comparing ***R*** to ***C***/2.

### Evaluation of CNV calling results on BCC samples

Both AtaCNV and Copy-scAT were run with default input and parameters. We transformed the outcome of Copy-scAT into a copy ratio matrix like AtaCNV, where rows and columns were bins and cells, respectively. To evaluate the outcomes of two methods, we obtained WES data of these BCC samples and non-tumor tissues from the previous study^21^. Then we applied BIC-seq2^17^ to call CNVs at bulk level. This algorithm divides genomes into bins of variable size that contain equal number of mappable positions and uses a regression model to estimate the expected read depth of each bin. Then, it infers breakpoints based on the observed read depth and the expected read depth. Specifically, we chose bin length of 1 million bp and tuning parameter of the segmentation step to be 5. We obtained the log2 copy ratio of each bin and denoted it by ***r***_*WES*_. From the single-cell profiles given by AtaCNV and Copy-scAT, we calculated pseudo-bulk CNV profiles using the log2 mean copy ratios of each bin over all tumor cells and denoted them as vector ***r***_*1*_ and ***r***_2_. Then, we evaluated the accuracies of the two methods by comparing ***r***_1_ and ***r***_2_ with ***r***_*WES*_. The overall similarity was measured by Pearson’s correlation between ***r***_*i*_ and ***r***_*WES*_. We also calculated bin-wise difference |***r***_*i*_ − ***r***_*WES*_| and plotted the box plot.

We also used circular binary segmentation (CBS) algorithm^42^ to call CNVs from WES datasets and evaluate the two methods using the CNVs inferred by CBS. In brief, we divided counts in tumor sample by counts in normal sample to obtain copy ratios in each bin and applied CBS algorithm to perform segmentation by R package DNAcopy^43^. We compared ***r***_1_ and ***r***_2_ with the new ***r***_*WES*_ likewise. The results are shown in Supplementary Figure 3F-G.

### Gene activity score from scATAC-seq dataset

We used ArchR’s function *addGeneScoreMatrix* to infer gene activity scores in every cells. The scores were calculated based on chromatin accessibility signals and could measure gene expressions when scRNA-seq data was not available. In the dataset PDAC 2, we used these scores to infer marker genes of each cluster using ArchR’s function *getMarkerFeatures* and assigned cell types based on marker genes. In the dataset aGBM 4218, we inferred marker genes for tumor subclones likewise.

### Tumor/normal cell classification analysis

#### Definition of CNV burden score

To discriminate malignant cells from non-malignant cells, we used the copy ratio matrix ***RR*** in the final segmentation to define CNV score and measure the CNV burden. We calculated the mean copy ratios of each bin over cells in the same cluster. The obtained *mm*-length vector ***r***^*k*^ (*k* = 1.2. …. *K*) represents the average copy state of the cells in cluster *k*. Then we used 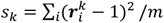 to measure the CNV burden of the cluster. Clusters with larger *s*_*k*_ are more likely to be malignant cells. Similarly, CNV scores for cells could also be defined using columns of ***R***. In each sample, we scaled the CNV scores of all cells by subtracting the mean and dividing the standard deviation, and then plotted the boxplots shown in Figure 2.

#### Tumor/normal classification from AtaCNV’s CNV profiles

We used the following method to classify tumor cells using CNV scores without the need for manually setting thresholds. We first obtained the smoothed count matrix ***X*** (as described in AtaCNV’s pipeline) and denoted the average counts of each bin in cluster *k* as a *m* -length vector ***x***^*k*^ (*k* = 1.2. …. *K*). Then, we measured the similarity between clusters by Pearson’s correlation and performed hierarchical clustering to obtain several groups (one group contained several cell clusters). We classified the group with the lowest average CNV score as normal cells and the others as tumor cells.

#### Tumor/normal classification from Copy-scAT’s CNV profiles

We transformed the outcome of Copy-scAT into a copy ratio matrix, then calculated CNV scores similarly. We assigned a threshold and classified clusters with CNV scores higher than the threshold as tumor cell clusters. This threshold was manually selected based on true labels, such that the classification results achieved the maximum accuracy.

#### Tumor/normal classification from predicted marker gene expression

We used ArchR to infer single-cell gene activity scores from scATAC-seq and used them to measure gene expressions. Then, we combined cancer type specific marker genes with UMAP plots to manually identify tumor cell clusters. The results were considered as true labels in this study.

#### Reference settings in different datasets

As described before, AtaCNV can use different information to infer a normal cell baseline. In the two PDAC datasets, we used paired normal samples. In EC and OC datasets, we randomly selected 50% of the true normal cells. In the four aGBM datasets and the two pGBM datasets, we did not provide normal cell information. As for Copy-scAT, we randomly selected 50% of the true normal cells in each dataset and used them as reference.

### Differential TF motif enrichment analysis

Activity scores of transcription factors were calculated using R package chromVar^44^ with the ENCODE^45^ motif database. We applied Wilcoxon’s rank sum test^40^ to detect differentially enriched TFs between two cell groups. P-values were adjusted by Benjamini-Hochberg’s method^46^. We also calculated the differences of average scores in two groups.

### Integration of scATAC-seq and scRNA-seq datasets in human gynecologic cancers

We used matched scRNA-seq data to better understand the transcriptional activity in human gynecologic scATAC-seq dataset. Firstly, we normalized the UMI count matrix of scRNA-seq and assigned cell clusters using Seurat. Then, we integrated scRNA-seq and scATAC-seq dataset using ArchR’s function (which was modified from Seurat’s function *FindTransferAnchors*), which measured the similarity between the gene activity score inferred from scATAC-seq and the gene expression from scRNA-seq. In brief, we could create a mapping between cell clusters of scATAC-seq and scRNA-seq by determining the cell having the highest similarity score in scRNA-seq data for each cell in scATAC-seq dataset. We used the matched scRNA-seq cell cluster for gene expression analysis of the scATAC-seq data.

### Cancer hallmark gene sets enrichment analysis

Using normalized gene expression profile from scRNA-seq dataset, we applied GSVA to calculate gene set enrichment score for 50 cancer hallmark gene sets from MSigDB^47^. We obtained 50 scores for each cell independently and applied Wilcoxon’s rank sum test to detect differentially enriched gene sets between two cell groups. P-values were adjusted by Benjamini-Hochberg’s method. In Figure 6C, the top 5 gene sets enriched in each group are displayed.

## Data availability

The scATAC-seq data used in this study are available in Gene Expression Omnibus (GEO) with the following accession code: BCC samples, GSE129785; PDAC samples, GSE147726; aGBM samples, GSE139136; pGBM samples, GSE163655; OC and EC, GSE173682. The 10X PBMC dataset is available at https://www.10xgenomics.com/resources/datasets/5-k-peripheral-blood-mononuclear-cells-pbm-cs-from-a-healthy-donor-1-standard-1-2-0. The paired WGS data of BCC samples are available in GEO with accession code PRJNA533341.

## Code availability

AtaCNV is available at https://github.com/XiDsLab/AtaCNV.

## Acknowledgements

This work was supported by the National Key R&D Program of China [2020YFE0204200 to R.X.], the National Natural Science Foundation of China [11471022, 71532001 to R.X.], China Postdoctoral Science Foundation [2022M720308 to Z.J.], and Sino-Russian Mathematics Center. Part of the analysis was performed on the High Performance Computing Platform of the Center for Life Sciences (Peking University).

## Author contributions

R.X. conceived and supervised the study. X.W. and Z.J. developed the model. X.W. and R.X. performed data analysis. R.X., X.W., Z.J., and Y.S. wrote the manuscript.

