## Supplementary Text for "Detecting copy number variations from single-cell chromatin sequencing data by AtaCNV"

**This supplementary file includes:**

Supplementary Figure 1 to 12

Supplementary Table1

### Supplementary Figures

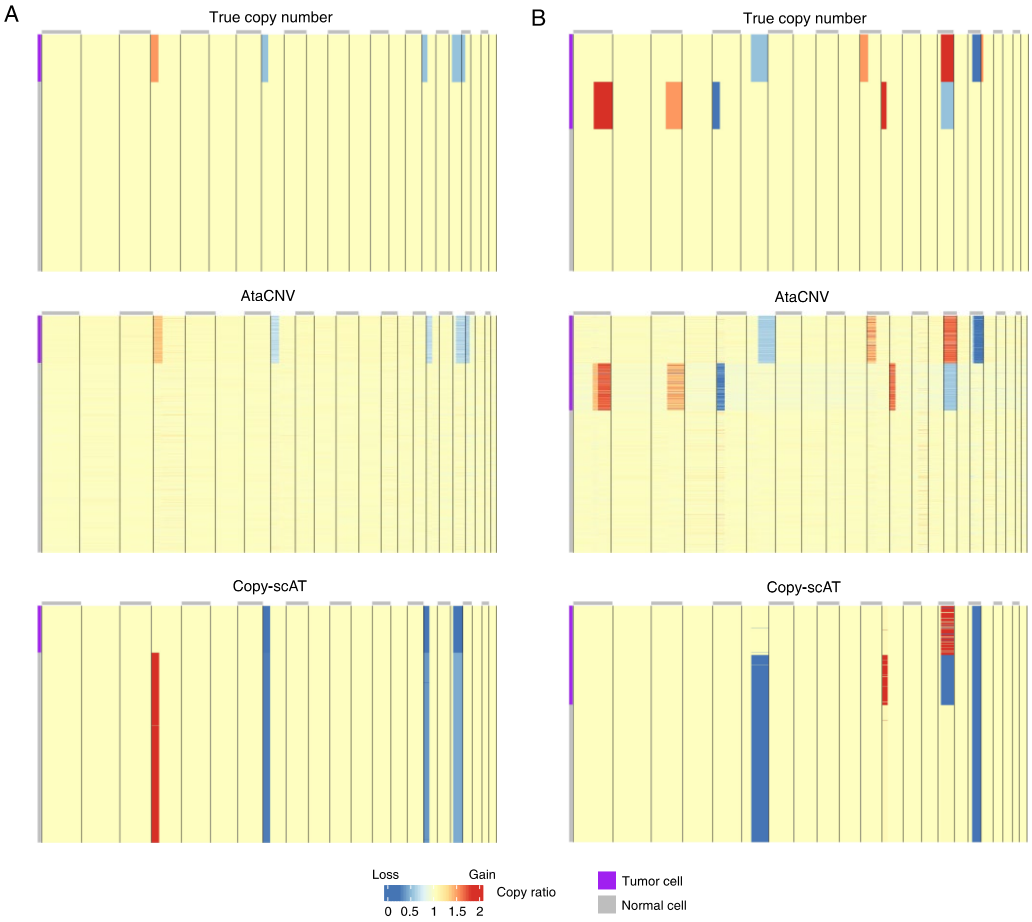


**Supplementary Figure 1. Examples of the simulation study. (A)** CNVs predicted by AtaCNV and Copy-scAT from a simulated dataset. The true CNV profile is shown at the top panel. **(B)** Ground truth and predicted CNVs from another simulated dataset. Copy-scAT incorrectly predicts CNVs in some chromosomal arms, leading to its lower precisions.

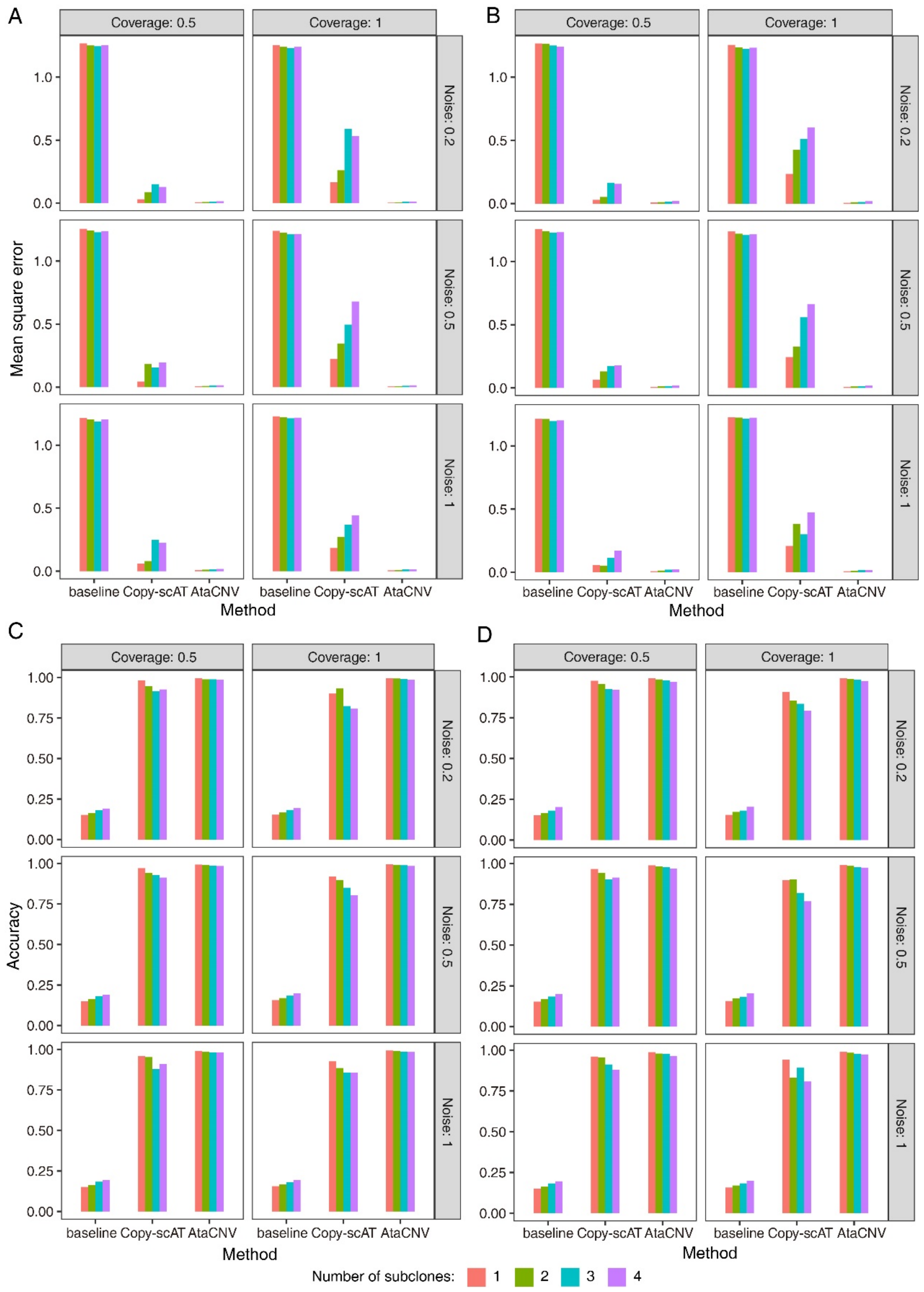


**Supplementary Figure 2. Performance of AtaCNV and Copy-scAT in the simulation study. (A, B)** The mean square errors (MSEs) of AtaCNV, Copy-scAT, and the baseline method in arm-level CNV simulation settings (A) and small scale CNV settings (B). **(C, D)** Similar to (A, B), but for the accuracies of different methods.

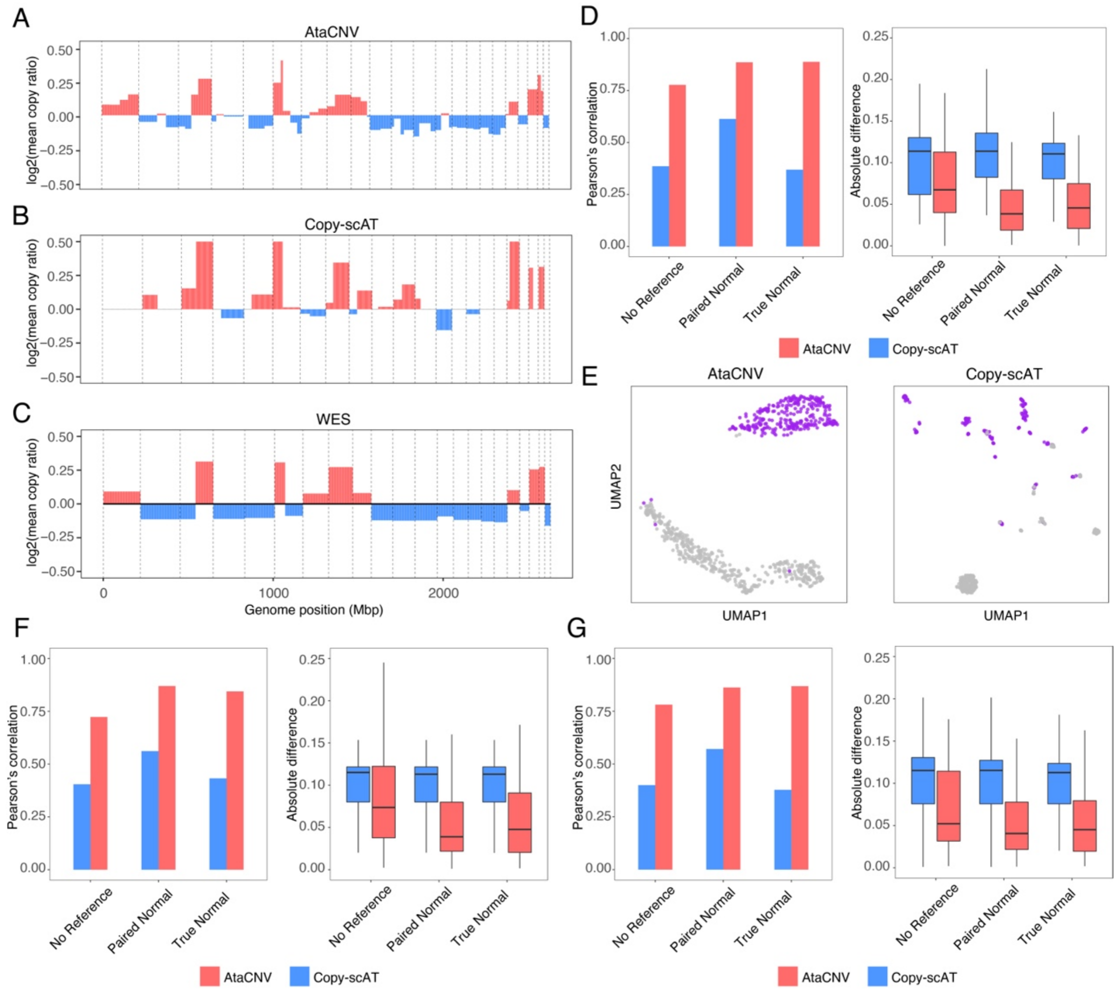


**Supplementary Figure 3. Evaluation of AtaCNV in the BCC samples. (A)** The mean copy ratio profile of another BCC sample from SU008 (pre-treatment) calculated by AtaCNV. **(B)** The mean copy ratio profile calculated by Copy-scAT. **(C)** The copy ratio profile inferred by matched WES dataset using BIC-seq2. **(D)** Comparison between WES result and mean copy ratios inferred by AtaCNV and Copy-scAT in terms of Pearson's correlation and relative differences. **(E)** The UMAP plots using CNV profiles from AtaCNV and Copy-scAT, colored by tumor/normal cell labels. **(F)** The performance of AtaCNV and Copy-scAT in the BCC sample from SU008 (post-treatment, the same sample shown in Figure 4), where WES CNVs were called by the circular binary segmentation (CBS) algorithm. **(G)** The performance of AtaCNV and Copy-scAT in the BCC sample from SU008 (pre-treatment), where WES CNVs were called by the CBS algorithm.

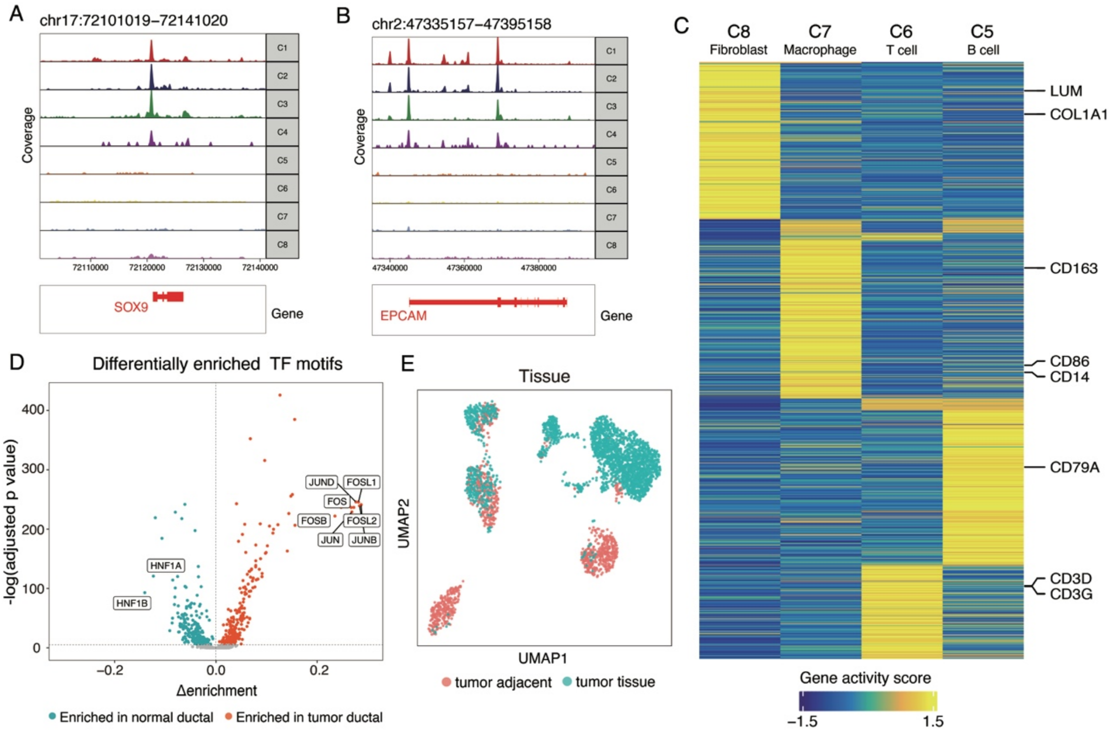


**Supplementary Figure 4. Cell type analysis of PDAC 2. (A, B)** Read depths of scATAC-seq data at ductal marker genes *SOX9* (A) and *EPCAM* (B) for different clusters. **(C)** Heatmap of marker genes' activity scores for different clusters. Known marker genes of B cell, T cell, macrophage, and fibroblast are annotated at right. **(D)** The differentially enriched TF motif test of C3 versus C1, C2, C4. From marker genes and TF motifs, C1, C2, and C4 are defined as tumor cell clusters. **(E)** The UMAP plot of peak signals colored by tissue types.

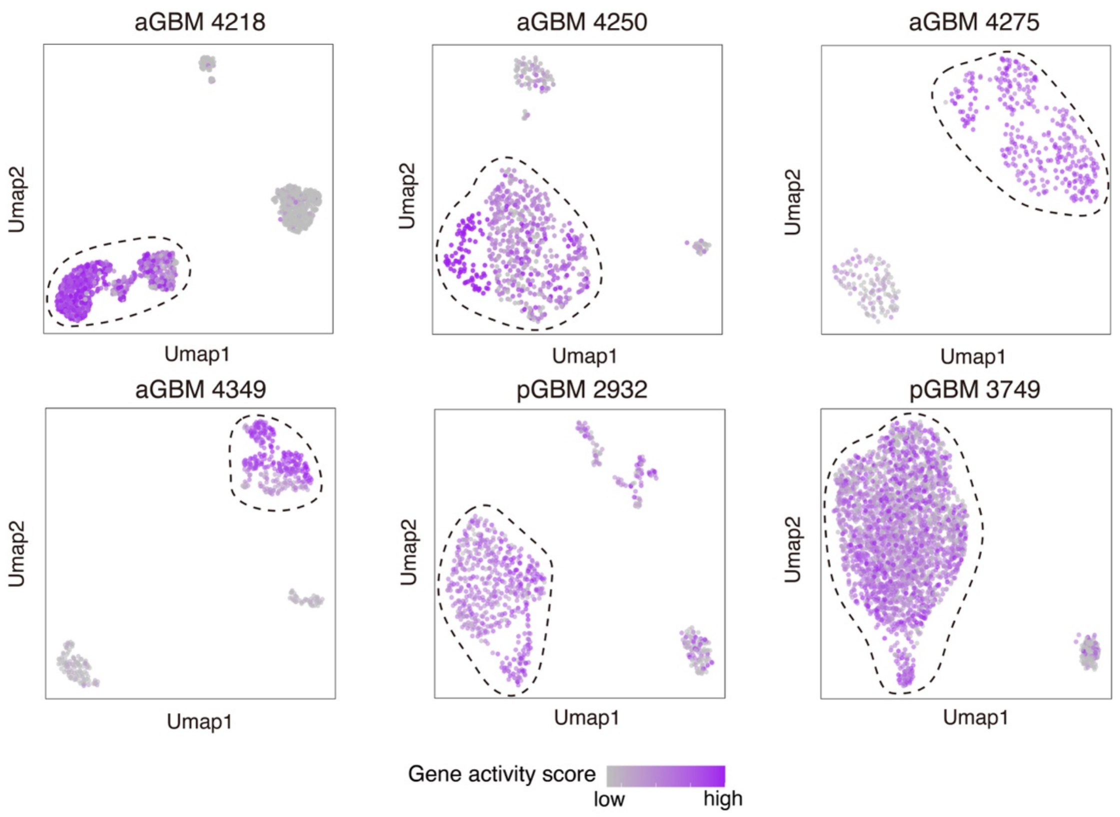


**Supplementary Figure 5. Cancer marker gene scores in GBM datasets.** The UMAP dimension reduction plots of cells in 4 aGBM and 2 pGBM datasets, colored by *EGFR* gene scores. Tumor cells identified by AtaCNV are circled in dash lines.

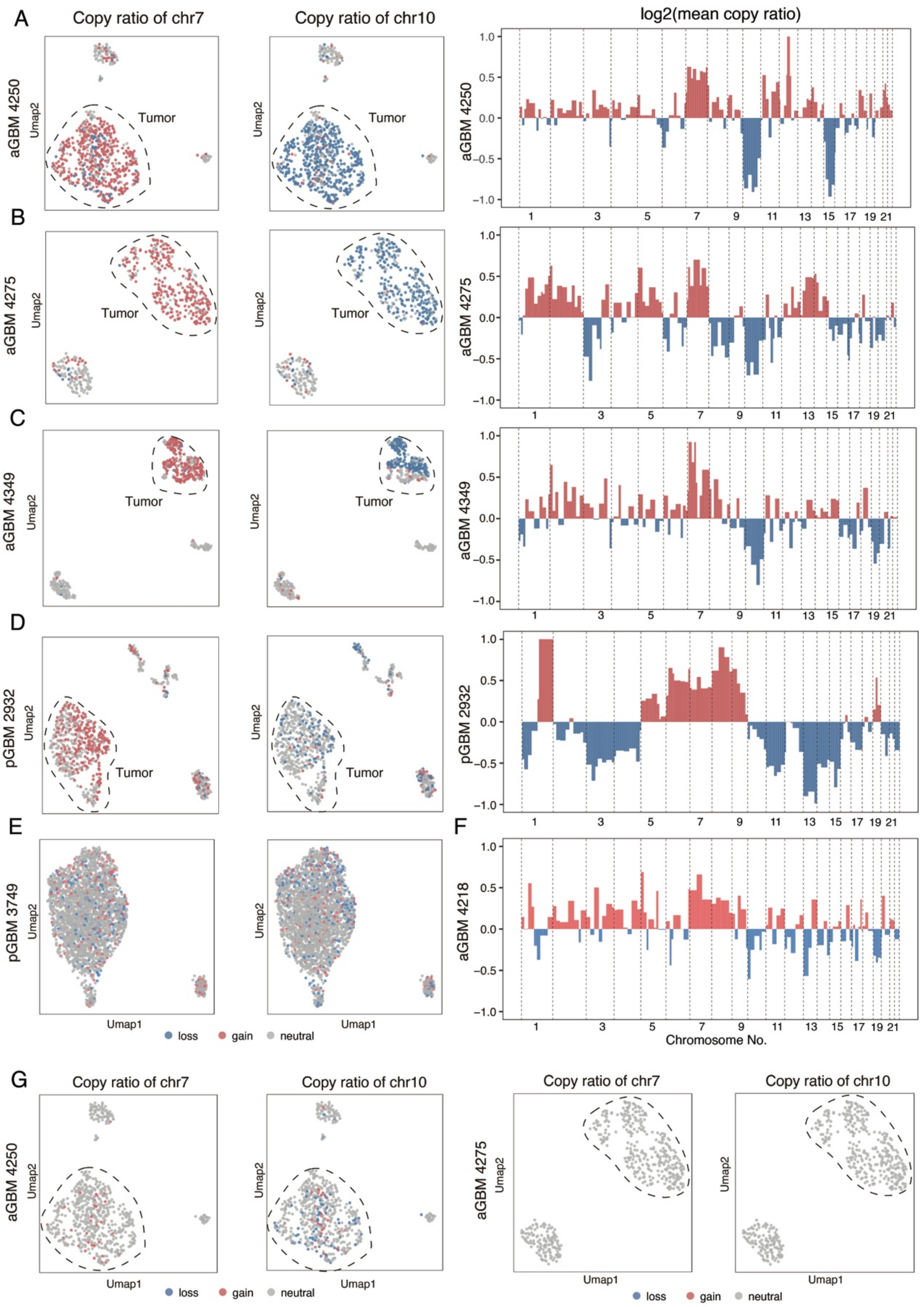


**Supplementary Figure 6. CNV profiles of GBM datasets. (A-F)** AtaCNV's results in aGBM 4250 (A), aGBM 4275 (B), aGBM 4349 (C), pGBM 2932 (D), pGBM 3749 (E), and aGBM4218 (F), respectively. The colors in the UMAP plots of peak signals represent the inferred copy number states by AtaCNV of chromosome 7 (left column) and chromosome 10 (middle column) in single cells. Tumor cells are circled in dash lines. The mean copy ratio profiles across tumor cells are shown on the right. Note that the corresponding UMAP plot of aGBM 4128 and mean copy ratio plot of pGBM 3749 are displayed in Figure 3H, I. **(G)** Copy-scAT's results of aGBM 4250 and aGBM 4275. The colors in the UMAP plots represent the inferred copy number states by Copy-scAT of chromosome 7 and 10.

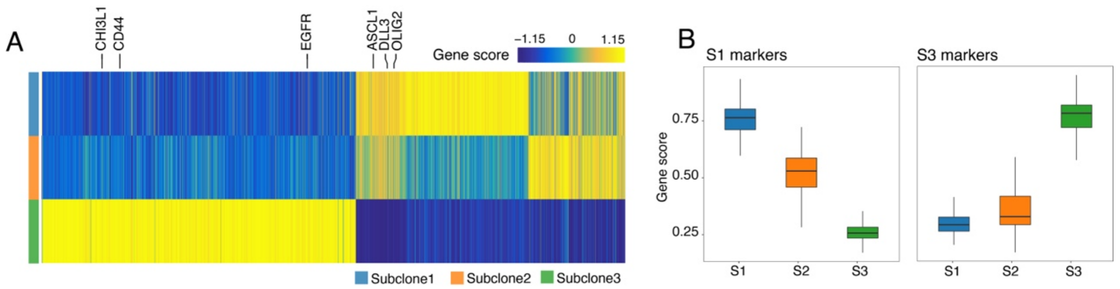


**Supplementary Figure 7. Subclone analysis of aGBM 4218. (A)** Heatmap of inferred marker genes' activity scores in different subclones. Known marker genes of PN subtype (*ASCL1*, *DLL3*, *OLIG2*) and MES subtype (*CHI3L1*, *CD44*, *EGFR*) are annotated at top. **(B)** Boxplots of S1 and S3 marker gene scores in three subclones.

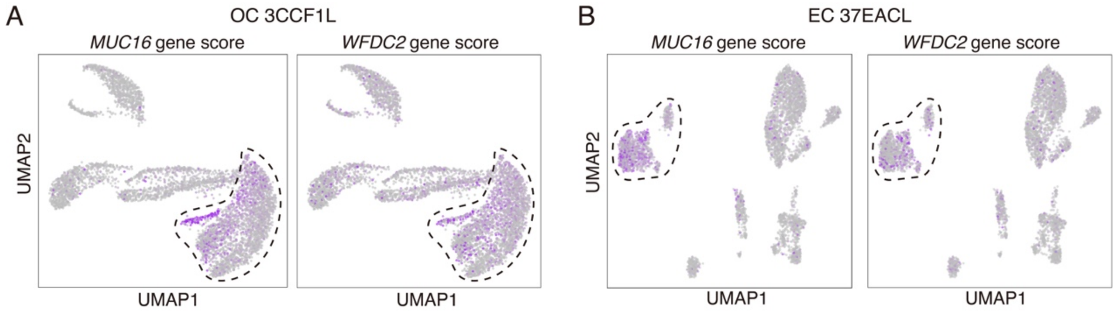


**Supplementary Figure 8. Cancer marker gene scores in the OC and EC datasets.** The UMAP plots of cells in OC (A) and EC (B) datasets, colored by *MUC16* and *WFDC2* gene scores. Tumor cells identified by AtaCNV are circled within dash lines.

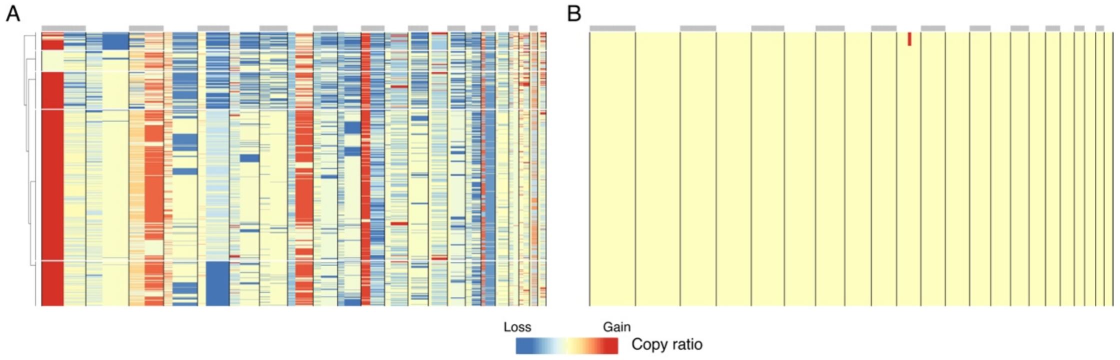


**Supplementary Figure 9. Copy-scAT's result in EC dataset. (A)** CNV detected by Copy-scAT in EC dataset. **(B)** Focal amplifications inferred by Copy-scAT.

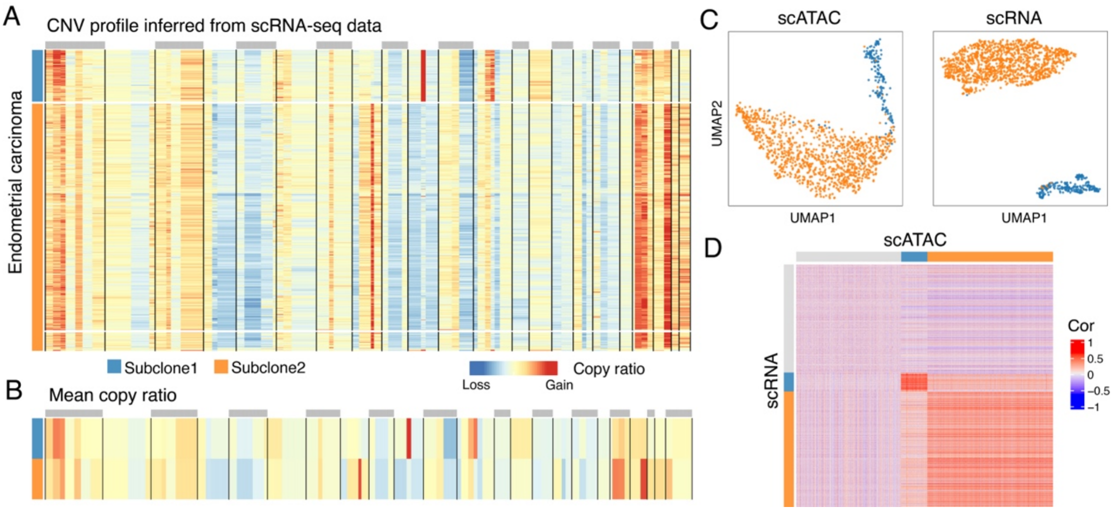


**Supplementary Figure 10. The CNV profile of the EC sample from scRNA-seq. (A)** Heatmap of single-cell copy ratios inferred from scRNA-seq data. Clustering analysis of the CNVs reveals two subclones. **(B)** Mean copy ratios of the two subclones. **(C)** The UMAP plot of tumor cells using CNV profile inferred from scATAC-seq (left) and scRNA-seq (right), respectively. **(D)** Heatmap of pairwise Pearson's correlations between the CNV profiles of cells inferred from the scRNA-seq and scATAC-seq dataset.

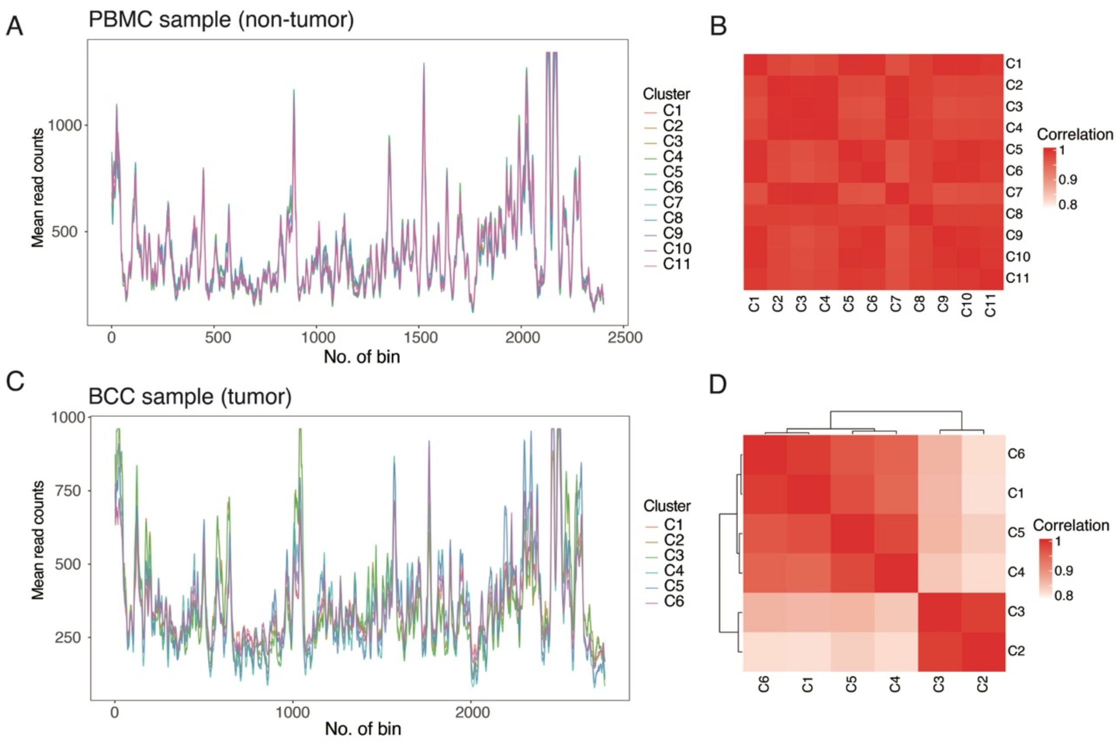


**Supplementary Figure 11. Similarities between cell types in normal and tumor samples. (A)** The average smoothed read counts of each cell type in the 10X PBMC dataset (non-tumor sample). **(B)** Pairwise Pearson's correlations of read counts between different cell types of the PBMC dataset. **(C)** The average smoothed read counts of each cell type in a BCC dataset (SU008 post, tumor sample). **(D)** Pairwise Pearson's correlations of read counts between different cell types of the PBMC dataset of the BCC dataset.

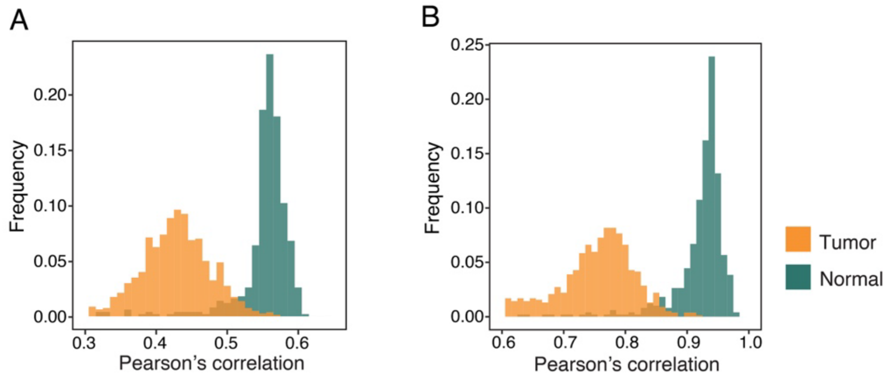


**Supplementary Figure 12. AtaCNV's normal cluster selection for the sample aGBM 4218**. **(A)** Histograms of Pearson's correlations of smoothed read counts with gene numbers in the bins for the known tumor and normal cells. **(B)** Histograms of Pearson's correlations of smoothed read counts with median smoothed read counts of normal cells from a different sample (aGBM 4250).

**Supplementary Table 1. scATAC-seq datasets information.**

The basic information for all the tumor scATAC-seq datasets involved in this study, including sample name, cancer type, total cell number, tumor cell number, average fragments per cell, and availability of paired non-tumor sample.

| Sample | Cancer type | Total cell | Tumor cell | Average fragments per cell | Paired non-tumor sample availability |
|---|---|---|---|---|---|
| SU008 pre | Basal cell carcinoma | 751 | 297 | 32805 | True |
| SU008 post | Basal cell carcinoma | 309 | 115 | 27761 | True |
| PDAC 1 | Pancreatic ductal adenocarcinoma | 3649 | 70 | 7261 | True |
| PDAC 2 | Pancreatic ductal adenocarcinoma | 3420 | 1696 | 11123 | True |
| aGBM 4218 | Adult glioblastoma | 1355 | 839 | 30822 | False |
| aGBM 4250 | Adult glioblastoma | 671 | 577 | 43426 | False |
| aGBM 4375 | Adult glioblastoma | 446 | 314 | 20200 | False |
| aGBM 4349 | Adult glioblastoma | 429 | 279 | 27395 | False |
| pGBM 2932 | Pediatric glioblastoma | 718 | 502 | 44843 | False |
| pGBM 3749 | Pediatric glioblastoma | 2765 | 2628 | 29098 | False |
| EC 37EACL | Endometrial carcinoma | 4718 | 1237 | 10651 | False |
| OC 3CCF1L | Ovarian carcinosarcoma | 5719 | 2878 | 14712 | False |
